# OTULIN Interactome Reveals Immune Response and Autophagy Associated with Tauopathy in a Mouse Model

**DOI:** 10.1101/2025.02.07.636114

**Authors:** Ling Li, Mingqi Li, Yuyang Zhou, David Kakhniashvili, Xusheng Wang, Francesca-Fang Liao

## Abstract

Tauopathies are neurodegenerative diseases that are pathologically characterized by accumulation of misfolded microtubule-associated protein tau aggregates in the brain. Deubiquitination, particularly by OTULIN, a unique deubiquitinase targeting methionine-1 (M1) linkages from linear ubiquitin chain assembly complex (LUBAC)), is reportedly associated with the accumulation of neurotoxic proteins in several neurodegenerative diseases, likely including tauopathies. To investigate the potential roles of OTULIN in tauopathies, we analyzed the OTULIN interactome in hippocampal tissues from PS19 transgenic (Tg) mice and their non-transgenic (nTg) littermate controls using affinity purification-mass spectrometry (AP-MS). We identified 705 and 800 proteins enriched in Tg and nTg samples, respectively, with a protein false discovery rate (FDR) of <1%. Of these, 189 and 205 proteins were classified as probable OTULIN interactors in Tg and nTg groups, respectively, based on Significance Analysis of INTeractome (SAINT) score of ≥0.80 and FDR of ≤ 5%. A total of 84 proteins were identified as OTULIN interactors in the PS19 Tg group, while 100 proteins were associated with OTULIN in the nTg controls. Functional enrichment analyses revealed that OTULIN-interacting proteins in the nTg group were enriched in pathways related to spliceosome, complement and coagulation cascades, and ribosome, whereas those in the Tg group were associated with immune response and autophagy. These findings suggest that OTULIN-interacting proteins may play a critical role in the pathogenesis of tauopathy in this mouse model.

**Highlights:**

1. OTULIN interactome in hippocampal tissues from PS19 transgenic (Tg) mice and their non-transgenic (nTg) littermate controls is analyzed.
2. OTULIN-interacting proteins in the nTg group are enriched in pathways related to spliceosome, complement and coagulation cascades, and ribosome.
3. OTULIN-interacting proteins in Tg are implicated in immune response and autophagy.
4. ATP2A2 is identified as an OTULIN-interacting protein specifically enhanced in Tg mice.

## Introduction

Tauopathies are a clinically and pathologically heterogeneous group of neurodegenerative diseases characterized by the abnormal accumulation of misfolded microtubule-associated protein tau aggregates in the brain [1]. This group primarily includes Alzheimer’s disease (AD), Pick disease, Frontotemporal lobar degeneration (FTD), Progressive supranuclear palsy (PSP), Corticobasal degeneration (CBD), and Chronic traumatic encephalopathy (CTE), each distinguished by unique patterns of tau aggregation and neurodegeneration. The mechanisms driving tauopathies are complex, involving genetic, biochemical, and environmental factors [2]. Understanding these mechanisms is crucial for elucidating disease pathogenesis and identifying effective treatments to halt or slow disease progression.

Mechanistically, misfolded tau aggregates are primarily degraded by the ubiquitin–proteasome system (UPS) and the autophagy-lysosome pathway (ALP) [3, 4]. Ubiquitination, one of the most pervasive post-translational protein modifications with ubiquitin, is widely recognized to serve as a targeting signal for protein degradation that is mediated by UPS or ALP [5]. Deubiquitination, the equally important reversal of the ubiquitination process, is mediated by deubiquitinases (DUBs). Recently, deubiquitination has gained attention as a potential therapeutic target in tauopathies including AD [6]. Emerging evidence suggests that dysregulation of deubiquitination can lead to protein aggregation and impaired protein turnover, both hallmarks of AD pathology [7].

OTULIN is the sole DUB known to specifically cleave Methionine-1 (M1) linkages mediated by the E3 ligase linear ubiquitin chain assembly complex (LUBAC) [8, 9]. The M1-linked ubiquitin chains are best characterized to regulate NF-κB activity, immune homeostasis, and responses to infection [10]. The ability of OTULIN to specifically target and cleave M1-linked ubiquitin chains renders it to precisely regulate NF-κB signaling pathways [11]. Aberrant NF-κB signaling has been implicated in neuroinflammation and neurodegeneration [12]. Therefore, understanding and modulating OTULIN activity could provide new insights and therapeutic approaches for neurodegeneration via fine-tuning NF-κB signaling and preventing neuroinflammatory responses.

Emerging evidence has linked linear ubiquitination to the degradation of misfolded proteins in neurodegenerative diseases. Proteins, such as Huntingtin, Optineurin, Ataxin-3, and TDP-43, were recently shown to be modified with polyubiquitin chains linked with M1 linkage (or linear linkage) [13]. Moreover, linear ubiquitination of Huntingtin and TDP-43 promoted their proteasomal degradation and decreased the intracellular aggregate formation [13]. Linear ubiquitin was also detected in the neurofibrillary tangles (NFT) from AD brains constituting intracellular aggregates of hyperphosphorylated tau protein [14].

To elucidate the potential roles of OTULIN in tauopathy-manifested neurodegeneration, we employed an Affinity Purification (AP)-based proteomic approach to profile the OTULIN interactome in the most commonly used mouse model of tauopathy (i.e., PS19) [15]. We chose to focus on the hippocampus of PS19 transgenic (Tg) mice and their non-transgenic (nTg) littermate controls at a symptomatic age. We performed a pull-down proteomic experiment using an OTULIN antibody, followed by quantification of the proteome using liquid chromatography-tandem mass spectrometry (LC-MS/MS). A comprehensive bioinformatics analysis was conducted to characterize the OTULIN interactomes and identified pathways enriched for OTULIN-interacting proteins in both nTg and Tg mice. We experimentally validated the OTULIN-ATP2A2 interaction, revealing an age-dependent alteration in PS19 Tg mice.

## Experimental Procedures

### Animal

All animal experiments were conducted under a protocol approved by the Institutional Animal Care and Use Committee at the University of Tennessee Health Science Center. PS19 mice, C57BL/6(B6);C3-Tg(Prnp-MAPT*P301S)PS19Vle/J, (stock #008169, Jackson Laboratory, Bar Harbor, ME), which is hemizygous for the P301S mutation in human tau gene, *MAPT*, driven by the prion protein (PrnP) promoter. The PS19 transgenic mouse line overexpresses 1N4R human tau with the frontotemporal dementia-associated P301S mutation [15]. PS19 mice and their littermate controls, aged 8 months (mixed sex), were used in this study. Three biological replicates were included. Mice were housed 3-4 per cage on a 12-hour light/dark cycle with ad libitum access to food and water.

### Affinity purification and protein pull-down

Potential interaction between OTULIN and targeted binding proteins was analyzed by protein pull-down assay. Hippocampal tissues (∼20 mg) from Tg and nTg mice were isolated and lysed by homemade IP lysis buffer (20 mM Tris (pH 7.0), 150 mM NaCl, 1 mM EDTA, 1 mM EGTA, 0.5% NP-40, 2 mM DTT, 0.5 mM PMSF, 20 mM β-glycerol phosphate, 1 mM sodium orthovanadate, 1 µg/ml leupeptin, 1 µg/ml aprotinin, 10 mM *p-*nitrophenyl phosphate, and 10 mM sodium fluoride) with protease inhibitor cocktail (04693116001, Roche). The lysates were purified by centrifugation at 13,500 rpm for 10 min and measured by Bradford protein assay (5000201, Bio-Rad). The same amount of protein (300 mg) was mixed with 30 ml dynabeads protein A (10002D, Thermo Scientific) and 5 ml OTULIN antibody (#14127, CST), which could be replaced by immunoglobulin gamma (IgG) (#2729, CST) as control groups to exclude the background signal and nonspecific association with antibodies or beads. The mixture was then supplemented with IP lysis buffer to a final volume of 1 ml and incubated on a tube rotator at 4°c overnight. The beads were gently washed with PBS three times for subsequent protein MS analysis. For co-IP gel electrophoresis, the beads were washed with IP lysis buffer three times and boiled in 2X b-mercaptoethanol-containing loading buffer (#1610747, BioRad) at 95°C for 10 min. All bait and IgG control experiments were conducted in triplicate.

### Sample Preparation for Mass spectrometry

Samples (magnetic beads with bound proteins) were transferred into 100 ml of digestion buffer (100 mM TEAB, pH 8.3) for processing. The estimated amount of protein per experimental sample was 2 mg. Sample proteins were reduced with 1mM DTT for 45 min at 50°c, alkylated with 5mM iodoacetamide for 20min at RT in dark, incubated with 5mM DTT at RT for 15min, and digested with 0.2 mg of trypsin/Lys-C protease mixture (V5073, Promega) overnight at 37°c shaking at 1100 rpm. Collected supernatants with peptide digests were vacuum dried.

### LC-MS/MS analysis

Each dried peptide sample was re-dissolved in 50 ml of loading buffer (3% acetonitrile, 0.05% TFA), and 5 ml was analyzed using LC-MS-MS method with 100 min LC gradient for peptide/protein identification and label-free quantification (LFQ). Raw MS data were acquired on an Orbitrap Fusion Lumos mass spectrometer (Thermo Fisher) operating in line with Ultimate 3000RSLCnano UHPLS system (Thermo Fisher). The peptides were trapped on an Acclaim PepMap 100 nanoViper column (75µm x 20mm, Thermo Fisher) at 5ul/min flow rate for 5 min. The trapped peptides were separated on an Acclaim PepMap RSLC nanoViper column (75µm x 500mm, C-18, 2µm, 100Å, Thermo Fisher) at 300nl/min flow rate and 40°c column temperature using water and acetonitrile with 0.1% formic acid as solvents A and B, respectively. The following multi-point linear gradient was applied: 3% B at 0-4 min, 5% B at 5 min, 25% B at 55 min, 30% B at 60 min, 90% B at 63-73 min, and 3% B at 76-100 min. Data dependent acquisition (DDA) method was used with 3 sec cycles and the following MS scan parameters. Full MS scans were performed in the Orbitrap analyzer at 120,000 (FWHM, at m/z=200) resolving power to determine the accurate masses (m/z) of peptides. The following data dependent MS2 analysis was performed on precursor ions with peptide-specific isotopic pattern, charge state 2-6, and intensity of at least 10,000. For MS2 scans, peptide ions were isolated using quadrupole isolation with 0.7 m/z window, fragmented (HCD, 30% NCE), and the fragment masses were determined in the Orbitrap analyzer at 30,000 (FWHM, at m/z=200) resolving power. Dynamic exclusion was applied for 30 sec.

### MS data analysis and peptide identification

The acquired raw MS data was analyzed by Proteome Discoverer 2.4 (Thermo Fisher) using Sequest HT search algorithm and human protein database (SwissProt, Mus musculus, TaxID 10090, v.2017-10-25). The reversed target database was used as decoy database. Full tryptic peptides were searched; 2 miss-cleavages were allowed. The searched fixed modifications included: carbamidomethylation of Cys residues. The variable modifications included oxidation of Met, acetylation of the protein N-terminus, and Met-loss modification of protein N-terminus. The precursor and fragment ion mass tolerances were set to 10 ppm and 0.02 Da, respectively. The raw data were filtered for the precursor ions with S/N of at least 1.5. The PSMs were filtered for further analysis using a delta Cn threshold of 0.05. The q-values were calculated at PSM level (Percolator), and then, at peptide level (Qvality algorithm) to control false discovery rate (FDR). The FDR threshold of 0.01 was used to validate and filter the data at PSMs and then at peptide levels. The validated/filtered peptides were used for the identification of the candidate precursor proteins. Candidate proteins with same set of assigned validated/filtered peptides were grouped. A candidate protein (or a group of proteins) was rejected, if none of the assigned validated/filtered peptides was unique to that protein (or to that group of proteins). Using Sum PEP protein scores, The Experimental q-values were calculated, and the retained candidate proteins were further validated (without filtering) using 0.01 (strict) and 0.05 (relaxed) FDR thresholds.

### Protein quantification

Peptide quantification was based on LC peak area with at least 5 data-points; the following parameters were used for feature detection, chromatographic alignment, and feature linking: 5 for minimal trace length, 10 ppm for mass tolerance, automatic for retention time (RT) tolerance, 5 for minimal signal-to-noise (S/N) threshold. Protein abundances were determined as summed abundances of assigned peptides. Unique and razor peptides were used for protein quantification. Log_2_ transformation of protein abundance was used for subsequent data analysis. After transformation, 0.01 was utilized for interpolating missing values.

### Analysis of interacting proteins with OTULIN

To calculate the probability of an identified protein being a true interactor in a purification sample compared to background contaminants, we used the protein expression values and Significance Analysis of Interactome (SAINT) version 2 [16–18]. Briefly, for each replicate (3 total replicates), bait protein pulldowns were performed as described above in batches alongside control purifications. Scores were generated to compare OTULIN pulldown samples with control samples.

### Principal-component analysis

Principal-component analysis (PCA) was used to visualize differences among samples. Intensities of top 100 variation proteins in all samples were used as features of PCA. The pairwise Euclidean distance between features was calculated. PCA was performed using the R package prcomp (version 3.4.0).

### Differential expression analysis

Differentially expressed proteins between the OTULIN pull-down group and the control were identified using the limma R package (version 3.46.0) [19]. The Benjamini-Hochberg (BH) method was used to control for multiple-testing. An adjusted *p*-value of 0.01 and a log₂ fold change exceedingly twice the standard deviation of control samples was used as cutoff values.

### Enrichment and network analysis

R package clusterProfiler (version 3.19) [20, 21] was used for enrichment analysis. Gene Ontology (GO) terms with a *p* value of 0.01 were defined as significantly enriched. The Metascape bioinformatics online tool (www.metascape.org) [22] was used to cluster enrichment results, with the following parameters: a minimum protein overlap of 3, an enrichment *p* value of 0.05, and a minimum 3 proteins per enriched group. Pathway visualizations were generated by Cytoscape 3.10.2 [23].

## Results

### OTULIN Interactome Profiled by LC-MS/MS

We used the PS19 transgenic mouse model, one of the most commonly used and well-characterized mouse models of tauopathy, which expresses the P301S mutant human tau associated with FTD and Parkinsonism linked to chromosome 17. As reported [15], tau aggregates develop at 6 months of age in these mice, and progressively accumulate in association with neuronal loss by 9-12 months of age. Neuronal loss and brain atrophy development starts off in the hippocampus and spreads to other regions including the neocortex by 12 months of age [15]. Moreover, prominent microglial activation has been detected long before tau aggregation. These mice also exhibit behavioral abnormalities recapitulating deficits seen in human tauopathies [24]. To investigate the role of OTULIN and its interacting protein partners in this tauopathy model, we conducted AP-based pull-down experiments on immunoprecipitated fractions from the hippocampal tissues (a brain region most critical for memory) of PS19 Tg mice and their nTg littermate control mice. Each condition was analyzed in three biological replicates, using an anti-OTULIN antibody and an IgG control antibody (Figure **1A**). In total, 12 samples were analyzed by LC-MS/MS. The raw MS data were searched against the mouse target-decoy database using the proteome discover (PD) program, resulting in the identification of 705 unique proteins in at least one of the 6 Tg samples, 800 unique proteins in at least one of the 6 control samples, with a protein false discovery rate (FDR) of 1%. Of these, 540 proteins were detected in both Tg and non-Tg samples, 165 proteins were detected only in transgenic mice, and 260 proteins were detected only in nTg mice. As expected, OTULIN expression levels were elevated in Tg samples compared to nTg samples (Figure **1B**). Within each group, the expression levels of OTULIN protein were enriched in the pull-down groups compared to the IgG control groups (Figure **1B**). Principal component analysis (PCA) demonstrated a clear separation between the OTULIN pull-down samples and control samples, as well as between Tg and nTg samples (Figure **1C**). This separation was further supported by sample-to-sample cluster analysis (Figure **1D**).

**FIGURE 1.**
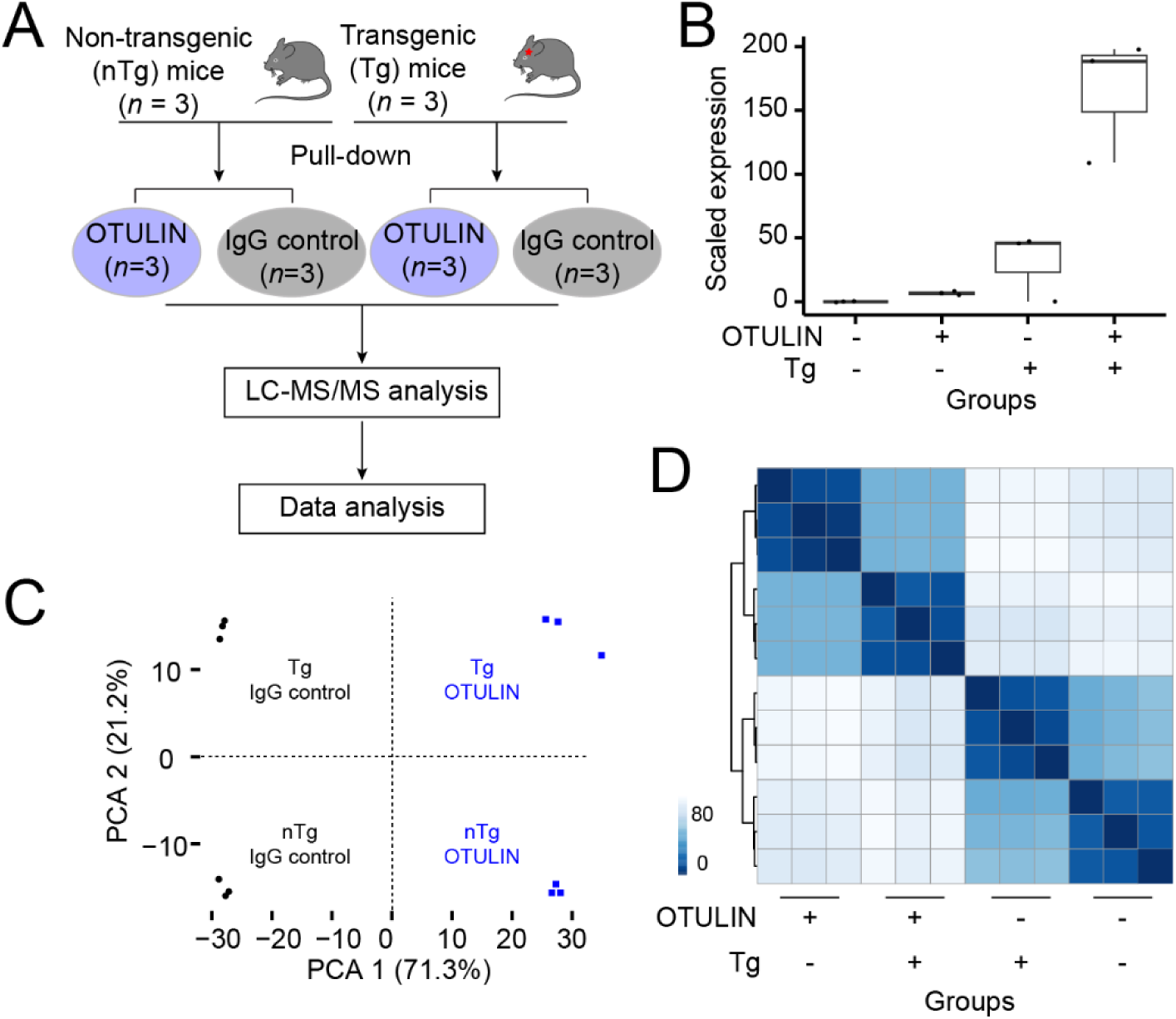
Workflow and quality control of the study. (A) Schematic diagram showing the experimental workflow. OTULIN pull-down experiments were conducted in both transgenic (Tg) and non-transgenic (nTg) mice, using IgG as a control. Samples were analyzed by liquid chromatography-tandem mass spectrometry (LC-MS/MS), followed by protein identification, and statistical analysis. (B) Bar plot comparing OTULIN protein expression levels across four conditions: Tg and nTg samples, as well as pull-down and IgG control groups. (C) Principal component analysis (PCA) plot showing distinct separation between Tg and nTg samples and between OTULIN and IgG control groups. (D) Heatmap illustrating sample-to-sample correlations, generated from the top 100 most variable proteins across 12 samples. The color intensity represents the correlation strength, with darker shades indicating stronger correlations between samples.

### OTULIN-interacting proteins in the nTg samples

To identify proteins interacted with OTULIN, we compared protein abundance in three samples pulled down by an OTULIN antibody with three matched IgG control samples in nTg. Using the limma R package [19], we identified 272 interacting proteins in nTg samples, with a 1% FDR and a fold change cutoff of 2.8 (Table **S1** and Figure **2A**). To further enhance confidence in these identified interactions, we calculated probability scores for protein-protein interactions using the Significance Analysis of INTeractome express (SAINTexpress) tool [17, 18]. SAINTexpress analysis detected 205 interacted proteins with SAINT confidence scores above 0.8 (Figure **2B**). Enrichment analysis revealed those OTULIN-interacted proteins enriched in the pathways, including mRNA progressing, ribosome, and U2-type spliceosome (Table **S2**; Figure **2C**). To examine those known OTULIN-interacted proteins, we mapped these 205 interacted proteins to protein-protein interaction (PPI) networks from the STRING database. We observed five proteins that interact with OTULIN, including TRIM21, MYCBP2, TAF, SKP1A, and USP9X. (Figure **2D**). Of note, among these, USP9X, an X-chromosome protein that escapes X-inactivation, was exclusively found to be interacted with OTULIN in the nTg samples, which was undetectable in the Tg samples. Prior studies revealed that lost or compromised function of USP9X leads to neurodevelopmental disorders in humans [25]. Potential implication of loss of the OTULIN-USP9X interaction in our symptomatic Tg samples in the development of tauopathy will be discussed later.

**FIGURE 2.**
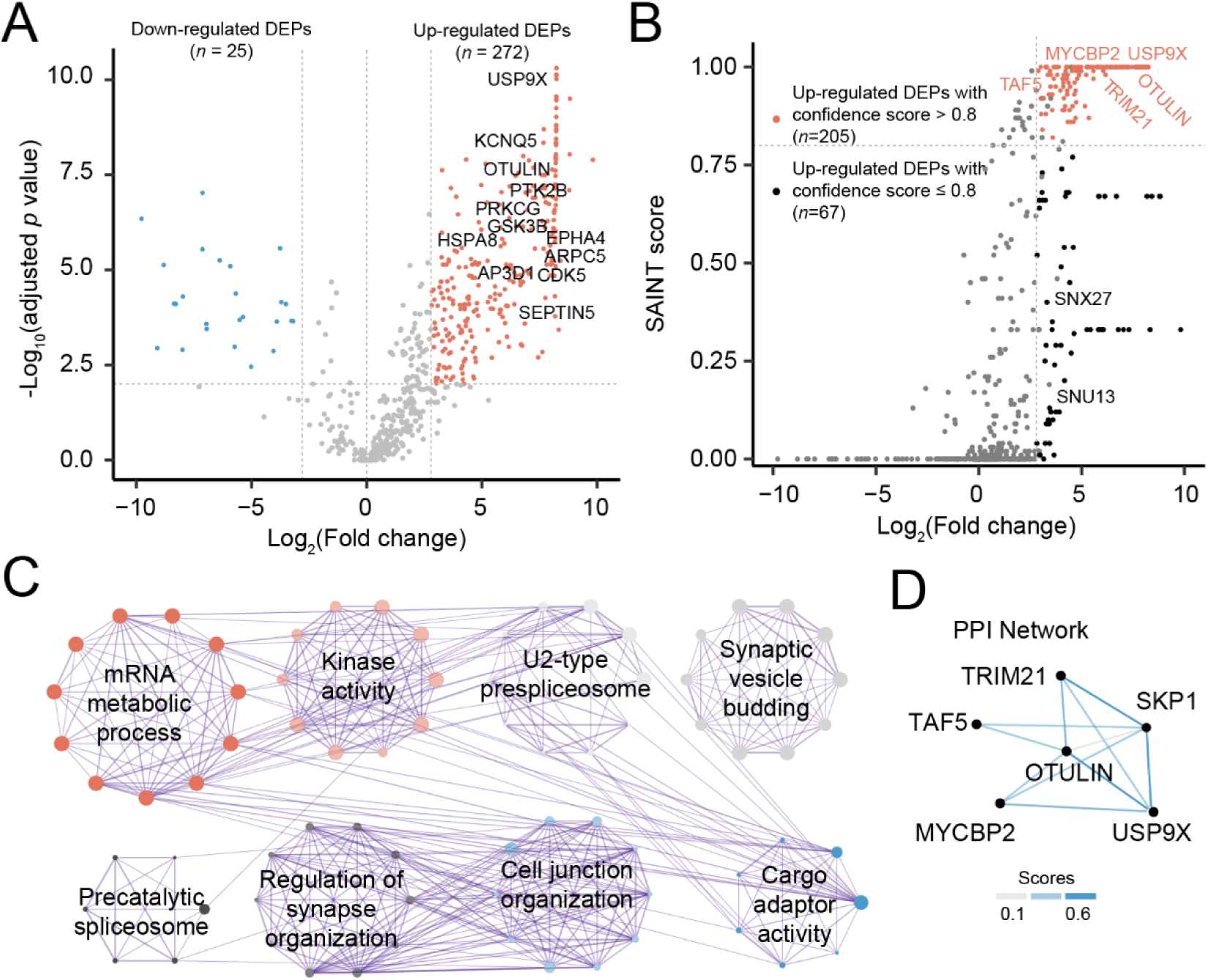
Interactomic analysis in the nTg mice. (A) Volcano plot showing differentially expressed proteins (DEPs) in the OTULIN pull-down samples compared to matched IgG control samples in nTg mice. The *x*-axis represents the log₂ fold change between pull-down and IgG control samples, and the *y*-axis shows the -log₁₀ (adjusted p-value). Proteins significantly upregulated in OTULIN pull-down samples (red dots) and downregulated proteins (blue dots) are highlighted. (B) SAINT score scatter plot displaying the OTULIN-interacting proteins in the nTg samples. The *x*-axis represents the log₂ fold change, while the *y-*axis displays SAINT probability scores, a statistical confidence metric for protein interactions. Proteins classified as high-confidence interactors (red) were defined by a SAINT score ≥ 0.8 and log₂ fold change ≥ 2.8. Several proteins, including OTULIN, are labeled for reference. (C) Metascape enrichment network visualization showing the top 8 clusters of functionally enriched GO clusters among OTULIN-interacting proteins. Nodes represent GO terms, with dot size proportional to the number of proteins in each pathway. Terms with shared cluster identity are positioned close to each other and colored accordingly. (D) Protein-protein interaction (PPI) network diagram showing OTULIN and its known interactors from the STRING database (https://string-db.org/). Line thickness corresponds to STRING confidence scores, with stronger interactions represented by thicker lines.

### Altered OTULIN-interacting proteins in the Tg tauopathy mice

To identify OTULIN-interacting proteins in PS19 Tg mice, we compared protein abundance in the three samples pulled down by the OTULIN antibody to the three matched IgG control samples in Tg mice. We identified 320 interacting proteins with a 1% FDR and a fold change cutoff of 2.1 (Table **S3** and Figure **3A**). Among these interacting proteins, SAINTexpress analysis revealed 214 proteins with SAINT scores above 0.8 (Figure **3B**). Enrichment analysis revealed those OTULIN-interacted proteins enriched in pathways, including autophagy and activation of immune response, in addition to some shared pathways enriched in nTg mice, such as mRNA processing, ribosome; and U2-type spliceosome (Table **S4** and Figure **3C**). For example, the immune response pathway includes several complement proteins (Figure **3D**), such as C1QA, C1QB, C1QC, C3, and C4B, which are key components of the complement system involved in innate immunity and inflammatory regulation.

**FIGURE 3.**
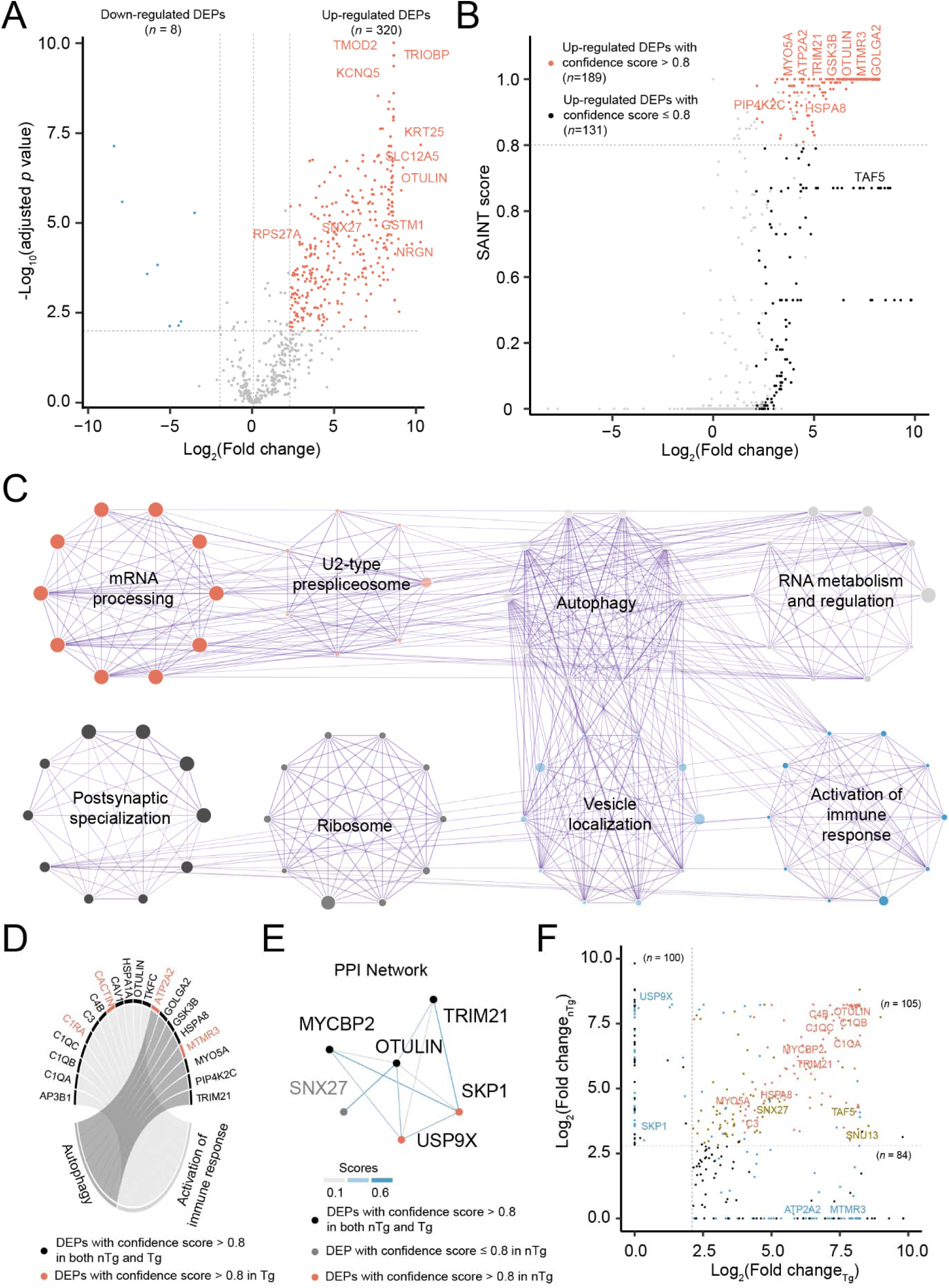
Interactomic analysis in the Tg mice. (A) Volcano plot displaying differentially expressed proteins (DEPs) identified in OTULIN pull-down samples compared to matched IgG control samples in Tg mice. The *x*-axis represents log₂ fold change, and the *y*-axis shows -log₁₀ (adjusted *p*-value). Proteins with significantly increased abundance in OTULIN pull-down samples are highlighted in red, marking 10 high-confidence OTULIN interactors. (B) Scatter plot of SAINT scores showing OTULIN-interacting proteins identified in Tg samples. The *x*-axis represents log₂ fold change, while the *y*-axis displays SAINT probability scores, a confidence metric for protein interactions. A SAINT score ≥ 0.8 and log₂ fold change ≥ 2.1 were used as cutoffs to define high-confidence interactors. Several key OTULIN interactors are marked. (C) Metascape enrichment network visualization showing the top 8 clusters of functionally enriched GO clusters associated with OTULIN-interacting proteins in Tg samples. Dot size reflects the number of proteins involved in each pathway. Nodes represent GO terms, with dot size proportional to the number of proteins involved in each pathway. Terms are color-coded by cluster identity, where functionally related terms are positioned close to each other. (D) Circos plot highlighting two representative enriched GO terms in Tg. The plot showing proteins associated with autophagy (left) and activation of immune response (right) pathways. Each node represents a protein, with lines connecting proteins and corresponding pathways. Red-labeled proteins were significantly altered only in Tg samples, while black-labeled proteins were significant in both nTg and Tg samples. (E) Protein-protein interaction network illustrating proteins interacting with OUTLIN, derived from the STRING database (https://string-db.org/). Line thickness corresponds to STRING confidence scores, with stronger interactions represented by thicker lines. Node colors denote significance levels under Tg or nTg conditions: black nodes are those OTULIN-interacting proteins with significant expression changes in both Tg and nTg. Grey nodes show the proteins with significant change at expression level, but no confidence score. Light red nodes are those proteins with signficant change at expression level and confidence scores, but only in nTg. (F) Scatter plot comparing altered OTULIN-interacting proteins between Tg and nTg mice, based on log_2_ fold change of expression levels and interaction significance scores. Red dots represent proteins with both significant expression changes and high-confidence interaction scores in both Tg and nTg samples. Golden dots indicate proteins with significant expression changes in both groups but without high-confidence scores. Light blue dots highlight proteins exhibiting either significant expression changes or high-confidence scores in only one condition (Tg or nTg). Black dots denote proteins with significant expression changes but lacking confidence scores in either group.

Mapping OTULIN interactors in Tg mice onto the protein-protein interaction network from the STRING database revealed the same four proteins (i.e., TRIM21, MYCBP2, and SNX27) identified in the nTg mice (Figure **3E**). The binding affinity between OTULIN and these partners are not significantly altered between nTg and Tg samples. Both TRIM21 and MYCBP2 are abundant forms of E3 ligases, although their roles in tauopathies remain obscure. Interestingly, GSK3β, a key signaling kinase crucial in metabolic regulation, was also detected to be interacting with OTULIN, but its interaction remained unchanged between the nTg and Tg samples. SNX27, a member of the sorting nexin (SNX) family of proteins that play a critical role in protein sorting and trafficking in the endocytosis pathway [26], was also identified. Of note, SNX27 is particularly enriched in the brain and has been recently reported as an OTULIN interactor [27], revealing a novel role of OTULIN in the regulation of endosome-to-plasma membrane recycling mediated by the major adaptor of the endosomal retromer complex (sorting nexin 27). SNX27 mediated retromer is known to be critical in regulating the tau clearance via endosomal-lysosomal degradation by autophagy mechanisms [28–30].

Comparisons of OTULIN-interacting proteins in Tg and nTg samples revealed 84 proteins only significantly in Tg samples, and 100 proteins only significantly in nTg samples, in addition to 105 proteins shared (Figure **3F**). For example, ATP2A2, an ATPase enzyme, was found to be interacted with OTULIN only in the Tg samples (*p* = 2.2 x 10^-22^). Of note, we identified two unique binding partners (i.e., USP9X and SPK1) with OTULIN only in the nTg samples, which were lost in the Tg samples. The potential gain or loss of function based on the selective binding of these molecules will be further discussed in the PS19 model (see the discussion section).

### The OTULIN-ATP2A2 interaction is altered age-dependently in PS19 Tg mice

Although the Ca^2+^ hypothesis was initially postulated for AD, it was later generalized to other neurodegenerative diseases including tauopathies [31]. Ca^2+^ uptake from cytosol to the endoplasmic reticulum (ER) plays an essential role supporting cellular Ca^2+^ homeostasis by maintaining a low cytosol Ca^2+^ concentration for proper neuronal signaling and activity [32, 33]. Accordingly, the Sarco/Endoplasmic Reticulum Ca^2+^ ATPase (SERCA) pump is identified to be the most major route to mediate Ca^2+^ entry from cytosol to ER and thus serves as such Ca^2+^-stabilizing mechanism for cellular Ca homeostasis. The SERCA (ATP2A) family consists of at least 12 isoforms thus far, deriving from tissue-dependent alternative splicing of the three genes [34]. SERCA 2 isoforms are predominantly expressed in the CNS with increasing importance in the neuronal calcium homeostasis [35]. ATP2A2 (i.e., SERCA2) is less characterized in the context of AD in terms of SERCA2a isoform than SERCA2b; the latter was dysregulated by presenilins PS1/PS2 mutants and associated with aberrant processing from amyloid precursor protein and with ER calcium overfilling in the context of AD [36]. More recently, ATP2A2/SERCA2 is reportedly linked to mental disorders, such as bipolar disorder and schizophrenia), suggesting its potentially important roles in brain functions [37].

SERCA is believed to play a significant role in the autophagy pathway by regulating intracellular calcium levels, where its activity is positively correlated with autophagic flux [38]. Lack of SERCA reportedly inhibits autophagosome-lysosome fusion, while SERCA activation increases autolysosome-like structures [39]. Given the critical role of autophagic dysfunction in the development of tauopathy [40], we further explored the potential functional importance of SERCA in the context of the OTULIN-SERCA axis in the PS19 model.

We first conducted additional experiments to verify the specifically enhanced interaction between OTULIN and ATP2A2 in the PS19 Tg samples by IP-Western blot analysis using a pan-antibody for SERCA2 (sc-376235). Indeed, we confirmed markedly enhanced OTULIN- SERCA2 interaction in late symptomatic Tg samples (10 months of age), while the direct input total SRECA2 protein abundancy was found to be reduced significantly in the Tg hippocampal samples at this late stage of tauopathy (Figure **4A**). Quantification based on densitometry analysis of this Western blot normalization of the OTULIN-pulled down SERCA2 against total SERCA2 protein expression levels revealed markedly enhanced physical interaction between OTULIN-SERCA2 in the Tg samples (Figure **4B**). We next compared the steady state SERCA2 protein levels in the Tg hippocampal samples at various ages and found that an age-related decrease in SERCA2 protein levels occurred starting from 4 months of age onward, and this decrease occurred in both the Tg and nTg hippocampi, while OTULIN expression remained unchanged for both genotypes at the two ages (Figure **4C**). We also determined the messenger RNA (mRNA) levels and found that ATP2A2 (SERCA2) mRNA levels decreased with age in both the Tg and nTg samples (Figure **4D, E**). Potential implication of the enhanced interaction between OTULIN- SERCA2 in the progression of tauopathy will be further discussed later (see the discussion section).

**FIGURE 4.**
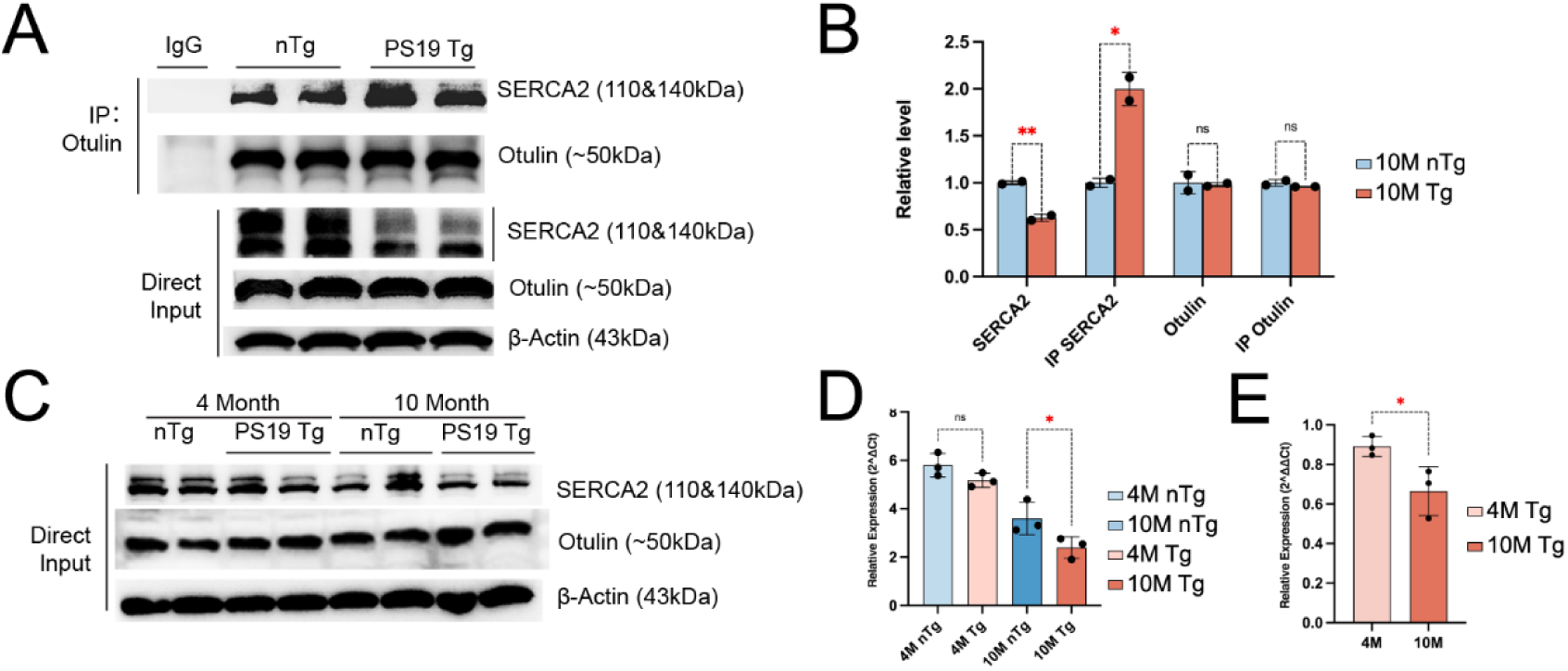
Enhanced OTULIN-SERCA2 interaction in symptomatic PS19 Tg samples. (A) Western blot analysis of SERCA2 and in hippocampal tissue lysates from OTULIN in 9–10-month-old (i.e., 10M) PS19 Tg and control (nTg) mice (*n*=2 mice per genotype). Upper panel: Immunoprecipitation (IP) was performed using an OTULIN antibody, followed by Western blotting to detect co-precipitated SERCA2 and OTULIN. Lower Panel: Direct input Western blot showing total protein levels of SERCA2 and OTULIN in whole-cell lysates, with β-Actin as a loading control. (B) Quantification by densitometry of the WB bands from Panel A. Multiple unpaired *t* tests (two tails?), \*\**p*_(SERCA2)_=0.006, \**p*_(IP SERCA2)_=0.016, *p*_(OTULIN)_ & *p*_(IP OTULIN)_= non-significant (ns). (C) Western blot detection of SERCA2, OTULIN in multiple age PS19 and nTg mouse brains. SERCA2 and OTULIN were detected in total cell lysates (Direct Input) and *β*-Actin was used as a loading control. (D) Relative ATP2A2 mRNA expression across different ages and genotypes. Bar plot shows ATP2A2 mRNA expression levels in 4-month-old (4M) and 10-month-old (10M) PS19 Tg and nTg mice (n=3 per genotype). One-way ANOVA comparisons test was used, *p*_(4M, nTg:Tg)_= non-significant (ns), \**p*_(10M, nTg:Tg)_=0.034. (E) Relative ATP2A2 mRNA expression ratio in PS19 Tg and nTg mice at the two ages. The bar graph shows the ratio of ATP2A2 (SERCA2) mRNA expression levels between PS19 Tg and nTg mice at 4 months (4M) and 10 months (10M) of age (n=3 per genotype). Statistical analysis was performed using an unpaired *t*-test (\**p* = 0.042).

## Discussion

Using advanced proteomic methodology, we separately analyzed the protein interactomes associated with OTULIN in symptomatic PS19 Tg and nTg mouse hippocampal tissue. For the first time, we identified different functional categories of OTULIN-interacting proteins in Tg compared to nTg brain tissue samples.

### Part I. OTULIN’s roles in immune responses

#### OTULIN -complement interactions

The complement system consists of over 30 complement factors under tight regulation. It is foundational to the innate immune response in defending the body against invading pathogens by phagocytosis or by the activation of the adaptive immune system [41]. In the CNS, the complement system protects the brain from not only pathogens but other potentially harmful stimuli such as aberrant proteins and cellular debris [42]. Hyperactivation of the complement system is associated with severe inflammatory responses in the CNS, companied by synapse and neuron loss in human AD, and also in tauopathy (e.g., FTD in the PS19 model), as best studied in the two seminal works [43, 44]: C1q - the initiating factor of the classical complement pathway, followed by activated C3-C3R [45]. All these components are elevated in human AD brains as well as in the late (9 months of age) PS19 Tg hippocampi.

We detected several complement components (C1QA, C1QB, C1QC, C3, and C4B) as high affinity binding partners of OTULIN in both nTg and Tg samples. Although the protein abundance of these OTULIN-interacted complements is not distinguishable between the nTg and Tg samples, given the widely reported upregulated complements in the symptomatic PS19 Tg hippocampal tissue at 9-months of age [43, 44], the actual pools of the OTULIN-coupled complements may be reduced during the pathogenesis in the PS19 Tg brains. Accordingly, we reason that OTULIN may be involved in modulating the stability of these major synaptic complements. OTULIN may also be richly localized in the post-synaptic density (PSD) compartment critical for maintaining post-synaptic homeostasis, a theory that warrants further investigation.

Among the complement system, we detected specific interaction between OTULIN-C1RA selectively in the PS19 Tg samples. C1RA or complement component 1 subcomponent r-A (C1ra), is a protein that plays a role in the complement system of the innate immune system. It is primarily secreted by hepatocytes, but is also expressed by other cells, including monocytes, possible macrophages and microglia, and endothelial cells. C1RA is an enzyme that activates C1s to its active form, which is the first step in the complement cascade involved in the innate immune processes (e.g., cell lysis and phagocytosis) [46]. The proteases C1r and C1s were initially discovered in a complex with C1q [47]. Of note, C1RA’s role in AD and tauopathy is not reported. Further studies of the interactions between OTULIN and these complements in the PS19 model will likely establish crucial regulatory roles and novel mechanisms in modulating multiple pathways in both innate and adaptive immunity in the CNS.

#### OTULIN-CACTIN interaction selectively detected in PS19 Tg samples

Toll-like receptors (TLRs) regulate gene expression by activating transcription factors, such as NF-κB and interferon-regulatory factors. NF-κB regulates multiple aspects of innate and adaptive immune functions, thereby serving as a pivotal mediator of inflammatory responses [48, 49]. Dysregulation of these pathways can lead to inflammatory diseases, and thus they are subjected to stringent control by negative regulators of innate immune signaling. Cactin (CACTIN) is a highly conserved eukaryotic protein among various organisms that was initially identified in Drosophila as an interactor of the “cactus” protein (i.e., the functional equivalent of IκB in mammals). The function of Cactin in the innate immunity has been studied from Drosophila to mammals, revealing that Cactin is a negative regulator of TLRs [50]. Recently, ubiquitin-specific protease 14 (Usp14) was identified as a major DUB for Cactin in *Drosophila* [51], and the specific member of the tripartite motif-containing proteins (TRIM39) functions as an E3 ubiquitin ligase for destabilizing Cactin to negatively regulate the NF-κB signaling induced by inflammatory stimulants such as TNFα[52]. Apart from being a negative regulator of the NF-kB signaling, Cactin is also part of the spliceosome and thus is involved in pre-mRNA splicing processes [53]. These two functions of Cactin implicate it being involved in several processes crucial in the CNS. However, its roles in brain have not been reported, nor in neurodegeneration including tauopathy. OTULIN’s best-characterized classic role is on negative regulation of NF-κB signaling [54, 55]. Further investigation to validate increased OTULIN-Cactin interaction in the PS19 model, in which overactivated NF-κB signaling plays a significant pathogenic role, will likely reveal novel regulatory roles of this conserved molecule in modulating innate immune response via Cactin-mediated mechanisms.

### Part II. OTULIN interactions with the ubiquitination system

#### OTULIN-USP9X interaction

We identified several OTULIN binding partners concerning protein ubiquitination (e. g., E3 ligases such as TRIM21 and MYCBP2). In particular, the ubiquitin-specific protease 9X (USP9X), a highly conserved member of the ubiquitin-specific proteases (USP) family of DUBs [25]. We found that USP9X interacts exclusively with OTULIN in nTg but not Tg samples in the PS19 mouse model. The implication of this loss of interaction in Tg mice is unclear. We speculate that this loss may contribute to Tau aggregation, leading to the disease pathogenesis in this tau model based on the known functional properties of these two molecules.

Both USP9X and OTULIN are DUBs with distinct yet complementary roles in ubiquitin signaling and thus are predictably involved in governing protein stability, cellular signaling, and immune responses. USP9X is one most diversified DUB, interacting with multiple E3 ligases [25, 56]. It plays a key role in stabilizing substrates such as MCL1, a pro-survival protein essential for neuronal development and synaptic plasticity [25]. In contrast, OTULIN is a specialized DUB that specifically cleaves linear (M1-linked) ubiquitin chains regulated by the linear ubiquitin assembly complex (LUBAC) [54, 55].

Dysregulation of USP9X has been linked to cancers and X-linked intellectual disability, as well as in CNS developmental disorders [25]. USP9X has been implicated in two neurodegenerative disorders: Parkinson (PD) and Diffuse Lewy Body Disease (DLBD). USP9X expression is altered in a mouse model of PD [56], and a portion of USP9X localizes to Lewy Bodies in PD and DLBD, modulating the stability of α-synuclein via autophagy-mediated degradation [57]. The functional implication of OTULIN-USP9X in nTg brain requires further investigation, along with the selective loss of this interaction in the symptomatic Tg brain may contribute to neurodegeneration related to tauopathy.

### Part III. OTULIN’s potential roles in autophagy

#### OTULIN-SNX27 interaction

The sorting nexin (SNX) family of proteins are involved in the trafficking of transmembrane proteins between the endosomal compartments, lysosome, trans-Golgi network, and plasma membrane. Sorting nexin 27 (SNX27), like VSP35, is a unique member of this family as it contains an additional PDZ domain in addition to a Phox domain. Evidence suggests that SNX27, in association with the retromer complex, binds cargo via its PDZ domain and recycles them from the early endosomes to the plasma membrane, thereby escaping their lysosomal degradation [26]. The PDZ binding motif is most commonly found in proteins involved in excitatory synapses, and thus located within regions of postsynaptic densities in the neurons. SNX27 is critical for postnatal growth and survival as SNX27-null mice die shortly after birth [58]. In the brain, SNX27 is found to be localized primarily within dendrites and has been shown to regulate synaptic plasticity[59, 60]. Therefore, dysregulated SNX27 functioning has been reported in several neurodegenerative diseases, such as AD. Dysregulation of SNX27, including its altered expression profile, has not been reported in terms of tauopathy. SNX27 knockout cells display increased autophagy due to impaired mTOR complex 1 (mTORC1) activation [61]. Interestingly, a recent study based on several cellular models of VPS35 knock out and overexpression discloses a potentially important role of VPS35, another PDZ-containing retromer like SNX27, in modulating tau clearance via the autophagy-lysosome axis [29] via their functional interaction between VPS35 and SNX27 [62]. Therefore, SNX27 likely plays an important role in autophagy, however, its involvement in this process is largely unknown.

OTULIN, in conjunction with LUBAC, reportedly regulate autophagy: *OTULIN* knockdown promotes autophagy initiation but blocks autophagy maturation to regulate the initiation and maturation of autophagy [63]. Given the pivotal role of APL pathway in tau clearance, the potential role of OTULIN in tau degradation warrants further investigation during the development of tauopathy. It is worth pointing out that physical and functional interaction and interplay has been validated in a recent work showing that SNX27-OTULIN binding, between their C-terminal PDZ-binding motif in OTULIN and the cargo-binding site in the PDZ domain of SNX27, antagonizes SNX27-dependent cargo loading, and thus the endosome to plasma membrane trafficking [27]. We anticipate this fundamental functional regulatory mechanism may be generally applied to most cell types, potentially crucial for autophagy and synaptic functions.

#### OTULIN -SERCA2 interaction

ER Ca^2+^ overfilling has been best studied in AD: ER calcium overfilling refers to an excessive accumulation of calcium within the endoplasmic reticulum (ER), which can lead to a subsequent “calcium overload” in the cytosol due to dysregulated release mechanisms, contributing to neuronal damage and disease progression; this is primarily caused by malfunctions in the presenilin proteins (PS) that act as ER calcium leak channels, leading to impaired calcium homeostasis within the ER. This theory was mostly based on prior research in cases of PS mutations/loss of function of presenilin (PS2 in particular) in ER, modulating ER calcium homeostasis. In most AD cases, especially those sporadic AD, it is unclear if the ER calcium dyshomeostasis is due to overfilling (insufficient SERCA activity) or upregulated ATP1 isoform (overactivated SERCA activity) ([64]. Indeed, several studies with pharmacological agents of SERCA modulators/activators successfully correct Ca uptake to ER and protect against ER-stress-induced neuronal death in cellular [65, 66] and mouse model [67, 68], supporting loss of functional mechanism of SERCA in AD. SERCA activity changes are not reported in tauopathy. Although ER stress is widely demonstrated in AD/tauopathy and in the PS19 model [68–70], the altered SERCA activity changes, as a result of ER-stress, are not reported in tauopathy.

Here in the PS19 model, we show loss of SERCA2 protein (steady state level by Western blot analysis) in an age-dependent manner, starting from 4 months of age. Based on the identified increased interaction between OTULIN and ATA2A2 (i.e., SERCA2) in the PS19 Tg hippocampal samples, we speculate the following theory: OTULIN as a unique DUB of linear ubiquitination by LUBAC may interact with ATP2A2 to mediate its deubiquitination, and thereby stabilizing it via preventing ubiquitination-induced protein degradation. This mechanistic hypothesis, along with the potential implication of the increased OTULIN-ATP2A2 in the ER-specialized functional processes (e.g., autophagy), both warrant further investigation. Interestingly, the recent report on the hyper-dopaminergic state detected in the brain specific in ATA2A haploinsufficient mice[37] due to prolonged cytosolic Ca^2+^ transients. Given the pivotal role of SERCA family members in ER Ca^2+^ homeostasis, and the ER stress in tauopathy development [69, 70], validation of the SERCA2 function in these aspects will facilitate therapeutic development in future.

### CRediT authorship contribution statement

Francesca-Fang Liao and Xusheng Wang designed research; Mingqi Li performed IP-pull down and sample preparation; David Kakhniashvili performed LC-MS; Ling Li conducted bioinformatic analysis; Yuyang Zhou performed biological validation; Ling Li, Xusheng Wang, and Francesca-Fang Liao wrote the paper.

## Acknowledgments

This work was supported in part by R01-AG072703 to F-F. L. and X. W. We thank Ryan Rager-Aguiar for assisting in manuscript proofreading.

## Data availability

The mass spectrometry proteomics data have been deposited to the ProteomeXchange

Consortium via the PRIDE [71] partner repository with the data set identifier PXD048286.

## Supplemetary Tables

Table S1: Differentially expressed proteins between OTULIN pull-down and matched IgG control samples in nTg mice.

Table S2.Enrichment analysis of differentially expressed proteins in nTg samples.

Table S3: Differentially expressed proteins between OTULIN pull-down and matched IgG control samples in Tg mice.

Table S4. Enrichment analysis of differentially expressed proteins in Tg samples.

